# Unveiling the anti-glioma potential of a marine derivative, Fucoidan: its synergistic cytotoxicity with Temozolomide-an *in vitro* and *in silico* experimental study

**DOI:** 10.1101/2022.12.02.518791

**Authors:** Indrani Biswas, Daisy S Precilla, Shreyas S Kuduvalli, Muralidharan Arumugam Ramachandran, S Akshaya, Venkat Raman, Dhamodharan Prabhu, T. S Anitha

## Abstract

**Introduction:** Glioma, coined as a “butterfly” tumor associated with a dismal prognosis. Marine algal compounds with the richest sources of bioactive components, act as significant anti-tumor therapeutics. However, there is a paucity of studies conducted on Fucoidan to enhance the anti-glioma efficacy of Temozolomide. Therefore, the present study aimed to evaluate the synergistic anti-proliferative, anti-inflammatory and pro-apoptotic effects of Fucoidan with Temozolomide in *in vitro* and *in silico* experimental setup.

**Methodology:** The anti-proliferative effects of Temozolomide and Fucoidan was evaluated on C6 glioma cells by MTT and migration assay. Modulation of inflammatory markers and apoptosis induction was affirmed at the morphological and transcriptional level, by dual staining and gene expression. Molecular docking (MD) and molecular dynamics simulation (MDS) studies were performed against the targets to rationalize the inhibitory effect.

**Results:** The dual-drug combination significantly reduced the cell viability and migration of glioma cells in a synergistic dose-dependent manner. At the molecular level, the dual-drug combination significantly down-regulated inflammatory genes with a concomitant upregulation of pro-apoptotic marker. In consensus with our in vitro findings, molecular docking and simulation studies revealed that the anti-tumor ligands: Temozolomide, Fucoidan with 5-(3-Methy1-trizeno)-imidazole-4-carboxamide (MTIC), and 4-amino-5-imidazole-carboxamide (AIC) had the potency to bind to the inflammatory proteins at their active sites, mediated by H-bonds and other non-covalent interactions.

**Discussion and Conclusion:** The dual-drug combinatorial treatment synergistically inhibited the proliferation, migration of glioma cells and promoted apoptosis; conversely with the down-regulation of inflammatory genes. However, pre-clinical experimental evidence is warranted for the possible translation of this combination.

## 1. Introduction

Glioma, a primary brain tumor arising from multi-step tumorigenesis of glial cells is often characterized by poor prognosis with a median 5-year survival rate of less than 10% [1]. The standard-of-care treatment for glioma comprises maximal surgical resection of the tumor mass (20 weeks of survival), followed by radiotherapy (36 weeks of survival by radiation and surgery) and chemotherapy (40-50 weeks survival) employing temozolomide (TMZ) [2]. Despite such multimodal approach, the prognosis remains dismal and tumor recurrence is inevitable. Thus, there is an urgent need for alternate therapies against glioma.

While the search for alternative chemotherapeutic agents to combat glioma aggressiveness is underway, it is not surprising that the traditional drug discovery pipeline is often hampered by high cost and low success burden. In this scenario, repurposing drugs would represent an attractive and effective strategy to improve the odds of new drug development success for glioma; As the safety and pharmacological profiles of the repurposed drugs are well-known, they can be streamlined for their passage through FDA more quickly, besides cost-reduction [3]. Interestingly, several repurposed drugs are currently under clinical investigation against glioma, thereby highlighting the efficiency of this systemic approach.

While choosing the potent drug candidate for repurposing, epidemiological reports have stated that combining natural compounds with conventional radio and chemotherapy enhances the sensitivity of the standard drug and alleviates therapy-associated complications in various cancers [4]. Among them, marine natural products (MNPs) have proven to be effective biological modulators. These MNPs have been predominantly noted for their potent biological activities such as antimicrobial, anti-infective, anti-inflammatory, or anti-tumor activity [5]. Thus, a plausible approach to utilize MNPs as a lead compound would accelerate the development of new anti-cancer drugs with increased efficiency and fewer side effects.

One such MNP is fucoidan (FU), a fucose-containing sulfated polysaccharide derived from brown seaweeds. The crude extracts of FU are commercially available as nutritional supplements in Asia and in some parts of Northern Europe. In recent times, the anti-tumor effect of FU has been studied intensively in colon[6], prostate, breast cancers,[7] and glioma[8]. Moreover, when FU was combined with cyclophosphamide, the combination had significantly inhibited metastasis in C57BL/6 Lewis lung adenocarcinoma mice model [9]. Nevertheless, the chemo preventive properties of FU in glioma have not been explored so far. Till date, FU has been known to enhance the epigenetic differentiation by restricting tumor growth in glioma cells as a monotherapeutic agent [10]. Though the effect of FU, as a monotherapeutic agent in glioma has been reported, their efficacy in synergism with TMZ as combined therapy and its underlying molecular mechanism has not been explored so far.

In this lacuna, the current study was designed to gain novel insights on the anti-tumor and pro-apoptotic efficacy of FU, both alone and in combination with TMZ against C6 glioma cells. Additionally, to predict and identify the mechanism of cytotoxicity induced by the ligands-TMZ and FU, molecular docking and simulation were performed.

## 2.0 Materials and Methods

### 2.1 Cell Culture and Chemicals

C6 rat glioma cell line (RRID: CVCL_0194; Passage number-48) and Human embryonic kidney cell line (HEK 293T; Passage number-10) was obtained from National Centre for Cell Sciences (NCCS), Pune, India. The cells were maintained at 37°C under 5% CO_2_ in nutrient mixture Ham’s F12 media and Dulbecco’s modified Eagle’s medium (DMEM), supplemented with 10% FBS and 1% antibiotic solution (100 U/ml penicillin and 100 μg/ml streptomycin). At the rate of 80% confluency, cells were seeded in 6-well, 24-well, or 96-well plates based on the experiments being performed.

Stock solutions of TMZ and FU were prepared in dimethyl sulfoxide (DMSO) and water. The aliquots were stored at -20ºC. On the day of the experiments, FU and TMZ were dissolved in the culture medium, in which the final concentration of DMSO was less than 0.1 % (v/v).

#### 2.1.1 Cytotoxicity Assay

The cytotoxicity of the drugs, either alone or in combination was assessed using MTT on C6 and HEK293T cell lines. Briefly, C6 and HEK293T cells were seeded in a 96-well plate at a density of 5×10^3^ cells/well and incubated for 24 hours. Cells were then treated with TMZ and FU at working concentrations of 10, 50, 100, 150, 200, 250, 300 µM. Following, cytotoxicity was assessed 24 hours post-treatment. Based on the IC_50_ values of the individual drug-treated cells, TMZ (IC_20_: 1.2 µM) was combined with varying concentrations of FU (10, 50, 100, 150, 200, 250, 300 µM). After 24 hours of dual-drug treatment, media was discarded completely and 20 µl of 5 mg/ml MTT was added to each well and incubated at 37°C for 3 hours. The purple-coloured formazan crystals formed were dissolved in 180 µl of DMSO. Absorbance was read at 570 nm spectrophotometrically (Molecular Devices Spectra-Max M5, USA). The absorbance value of untreated control cells was fixed as 100%. The experiment was repeated at least three times. Cytotoxicity was evaluated using the following formula:

Cytotoxicity = [(Optical density {OD} of treated cell − OD of blank)/ (OD of control − OD of blank) × 100].

#### 2.1.2 Calculation of Selectivity Index (SI)

To determine the selectivity and specificity of TMZ, FU, and their combination on glioma cells, SI was calculated using the following formula: SI= (IC_50_ of the individual and dual-drugs on HEK293T)/ (IC_50_ values obtained for the same treatment regimen on C6 glioma cells). A higher SI value indicates more selective and sensitivity towards cancer cells of the probed drugs and their combination.

#### 2.1.3 Combination index analysis, normalized isobologram, and Fa-CI plot

The statistical combinatorial drug Index (CI) was calculated to analyse the interaction pattern of the drugs, while the Chou-Talalay method was used to determine the CI value. The CI values were calculated manually using the CI equation: CI=(D)1/(Dx)1+(D)2/(Dx)2, where (Dx)1 and (Dx)2 represents the dose of drug 1 and drug 2 in a combination which is required to attain same efficacy as that of drug 1 (D1) and drug 2 (D2), when used alone. According to the Chou-Talalay method, the following criteria of drug-drug interactions were considered by CI: synergistic effect, CI < 1; additive effect, CI = 1; antagonist effect of drugs, CI > 1. Further, Dose-effect curve, CI, Fa-CI and Drug receptor index (DRI) plots were generated using Compusyn.

#### 2.1.4 Wound-healing assay

To determine the effect of the drugs on the inhibition of glioma migration, C6 cells were plated at a density of 4×10^5^ in 6-well plates. At 90% confluency, using a 200 µl sterile plastic tip, a scratch was made on the monolayer of cells. The peeled-off cells were washed using PBS and thereafter the cells were maintained in a 2% FBS medium. Following, the cells were treated with the indicated concentration of the drugs both individually and in combination for 24 hours. Cell scratch spacing was evaluated at 0 and 24 hours by calculating the cell mobility[11].

#### 2.1.5 Haematoxylin and Eosin (H&E) Staining

To detect the induction of apoptotic changes at the morphological level, H & E staining was carried out[12]. For this, 3×10^5^ of C6 cells were seeded in sterile coverslips in a 30 mm dish. Following 24 hours, cells were treated with the respective IC_50_ values of the drugs for 24 hours. Post incubation, the medium in each dish was discarded, followed by fixation of cells using 4% paraformaldehyde for 10 mins at room temperature. H & E stain was then performed and the cells were imaged at 40x magnification under a light microscope (Zeiss Light Microscope, Germany) and quantified using Image J.

#### 2.1.6 Acridine orange/ ethidium bromide (AO/EtBr) dual staining

To further, validate the changes associated with apoptosis at the morphological level, acridine orange/ ethidium bromide (AO/EtBr) dual staining was performed. C6 cells were seeded in a 24-well plate at a density of 20×10^3^/well and were then treated with the drugs, both individually and in combination for 24 hours at 37°C. Following incubation, cells were washed with PBS and stained with 5 μl of 100 μg/ml AO and 100 μg/ml EtBr at room temperature for 5 minutes. Stained cells were subsequently observed and imaged under a fluorescent microscope (Axiovert 40 CFL, Germany) with an excitation wavelength of 300-360 nm. The number of apoptotic cells per field was counted, and the apoptosis rate was calculated as: % apoptotic cells = number of apoptotic cells/total number of cells ×100, following quantification using ImageJ software.

#### 2.1.7 Nuclear Staining

To synchronously confirm the apoptosis induction effect of the drugs at the nuclear level, the fluorescent nuclear stain, 4’6, Diamidino-2-Phenyl Indole (DAPI) was employed. To carry out DAPI staining, C6 glioma cells were seeded at a density of 15×10^3^ cells/well and treated with the respective IC_50_ value of the drugs. After 24 hours post-treatment, the cells were washed with 1X PBS followed by fixation in 4% paraformaldehyde for about 10 minutes at room temperature. After fixation, the cells were permeabilized using Triton X-100 followed by incubation with 1μg/ml of DAPI in dark. Stained cells were subsequently observed and imaged under a fluorescent microscope (Axiovert 40 CFL, Germany) with an excitation wavelength of 359 nm. The percentage of apoptosis was evaluated using ImageJ software.

#### 2.1.8 Quantitative Real-Time PCR

To quantitatively assess the mRNA transcript levels of inflammatory and pro-apoptotic markers, quantitative Real-Time PCR (qRT-PCR) was performed. Briefly, total RNA from all of the experimental groups was isolated using RNAiso Plus (Takara, Japan). Complementary DNA (cDNA) was synthesized with HiMedia-cDNA Synthesis Kit (HiMedia, India). qRT-PCR was performed using CFX 96 thermocycler (Bio-Rad, Hercules, CA) and SYBR Green (Takara, Japan) to detect the mRNA. The reaction conditions were as follows: Initial denaturation: 95°C for 30 seconds; followed by 39 cycles of denaturation at 95°C for 10 seconds; annealing at a primer specific annealing temperature for 30 seconds. The relative gene expression was calculated with the 2^-∆∆C^_t_ method, where ∆∆C_t_ = (C_t target gene_ - C_t GAPDH_) sample- (Sample C_t target gene_ – Control C_t target gene_) calibrator. The primers were designed using the primer BLAST tool and are listed in Table 1.

**Table 1:**
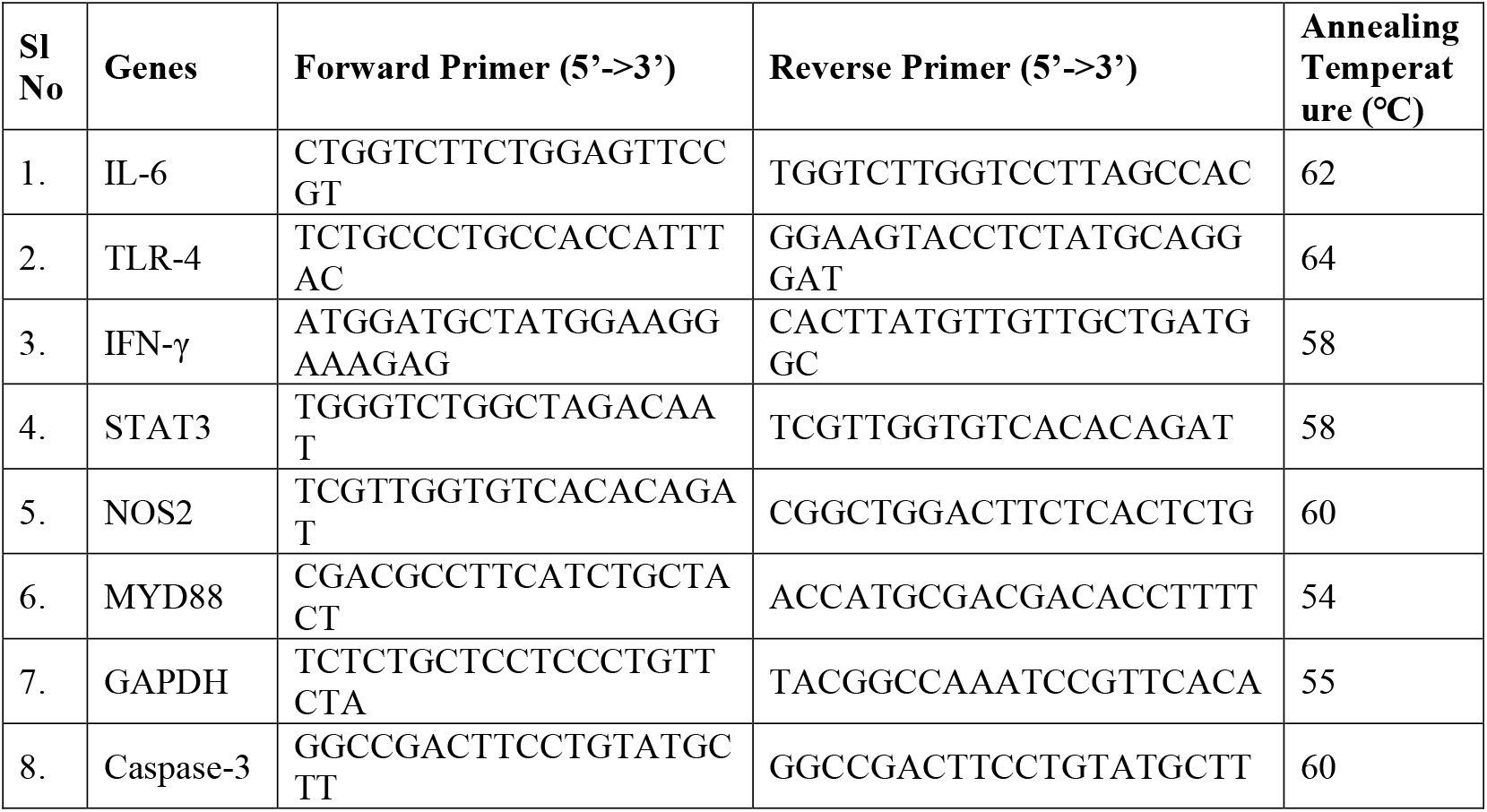
Primers designed for each gene

### 2.2 Molecular Docking

#### 2.2.1 Target retrieval

Target retrieval was performed for six major proteins of inflammatory pathway namely IL-6, TLR-4, JAK2, STAT3, IFN-γ, and NOS2 from UniprotKB (http://ca.expasy.org/sprot/). The 3D structure obtained from PDB was used for further analysis. Detailed information about the targets are listed in Table 2.

**Table 2:** List of targets retrieved from Uniprot and PDB

| Sl. No | Protein Name | Function | PDB ID | Structure Resolution (Å) |
| --- | --- | --- | --- | --- |
| 1. | IL-6 | Pro-inflammatory cytokine involved in innate immune response. | 2L3Y | NA |
| 2. | TLR-4 | Cooperates with lipopolysaccharide (LPS) to mediate the innate immune response. | 2Z64 | 2.84 |
| 3. | IFN-γ | Belongs to class of interferon family, aids in immunogenic response against foreign antigens, thereby activating effector immune cells and enhancing antigenic presentation. | 1FYH | 2.04 |
| 4. | JAK2 | Involved in various processes such as cell growth, development, and differentiation or histone modifications. Also involved in innate and adaptive immunity. | 2HDX | 2.35 |
| 5. | STAT3 | Mediates cellular responses to interleukins and also acts as a regulator of inflammatory responses. | 1BG1 | 2.25 |
| 6. | NOS2 | Messenger molecule with diverse functions, such as generation of nitric oxide and serves as a secondary messenger. It can mediate tumoricidal activity in macrophages. | 3NQS | 2.20 |

#### 2.2.2 Target preparation and active site prediction

The 3D structure of six targets was imported individually into the maestro and was prepared using protein preparation wizard of the Schrodinger suite (Schrödinger Release 2019–4: Maestro, Schrödinger, LLC, New York, NY, 2019). The receptor preparation process includes: modelling of missing residues and missing side chains, the addition of missing hydrogen atoms, correcting the missing atoms in residues, redundancy validation in occupancies of atoms, modification of metal ionization states and assigning proper charges, removal of undesired water molecules and ideal protonation state of histidine. The presence of negatively and positively charged amino acid residues facilitates to interact with the surrounding moieties across the ligand-protein complex, thereby rationalizing the presence of these three residues specifically, namely Arg, Gln and His. These amino acid residues of the drug targets were potentially spun to facilitate hydrogen and other non-bonded interactions, across the ligand-protein complexes with the surrounding moieties. The receptors were optimized using OPLS-3e force field and subsequently minimized to relax the bond angles, lengths and associated clashes. After preparation for all the six targets, the binding site information, which is used for the prediction of protein-ligand interactions, was collected using sitemap protocol (Schrodinger suite).

#### 2.2.3 Ligand retrieval and Preparation

The ligands selected for this study were TMZ, FU from *Fucus vesiculosus*, 5-(3-Methy1-trizeno)-imidazole-4-carboxamide (MTIC) and 4-amino-5-imidazole-caroboxamide (AIC). TMZ being a pro-drug, at physiological pH is rapidly hydrolyzed into the active form of the drug, 5- (3-methyltriazen-1-yl) imidazole-4-carboxamide (MTIC). This subsequently gets metabolized to produce the final degradation product of TMZ, 4-amino-5-imidazole-carboxamide (AIC). The 3D structures of ligands, TMZ, FU, MTIC and AIC were retrieved from Pubchem. The retrieved files were converted into .pdb format using Pymol. All the compounds were imported individually into maestro and were prepared using LigPrep module (Schrödinger Release 2019–4: Maestro, Schrödinger, LLC, New York, NY, 2019).

#### 2.2.4 Molecular docking analysis

The Glide XP (extra precision) docking method was used to predict the binding effectiveness of compounds in the active site of targets (Schrodinger suite) using Sitemap protocol as listed in Table 1. The prepared targets and ligands were given as input for glide docking and the grid box was produced utilising the receptor grid generation procedure (Schrodinger suite) to delineate the docking area in the protein, which allows for more accurate ligand binding scoring. The simulations of molecular docking were run with default parameters[13]. The docking score, glide energy and number of hydrogen bond (HB) contacts between the target and the ligand were used to validate the simulation results.

### 2.3 Molecular Dynamics Simulation (MDS)

The structural and dynamic changes accompanied with each of the protein-ligand complexes were analysed using molecular dynamic simulation. The simulation was executed with Gromacs 5.1.4 suite with GROMOS96 43a1 force field on LINUX based workstation. Gromacs provides high molecular simulation results and has been considered as a standard reference tool for simulation in comparison to Desmond. During the initial stage, system relaxation was performed to eliminate unwanted atomic contacts, which may lead to unstable MD simulation. Each of the protein-ligand complexes was solvated in a cubic simulation box under a Simple Point Charge 216 (SPC 216) water environment. There was a free run for 10 ps under the equilibrium period with a force constant of 1000 kJ mol−1 mol−2. Following energy minimization, the system was equilibrated with reference temperature of 300K at constant temperature and volume (NVT). Further, pressure was maintained with reference pressure of 1 bar at constant temperature and constant pressure (NPT). Temperature was controlled by V-rescale, a modified approach for Berendsen temperature coupling method. The minimized structures were subjected to MD simulation for 20000 picoseconds (ps) without restrictions. Analysis for MD simulation was executed in terms of Root-Mean-Square-Deviation (RMSD), Root- Mean- Square- Fluctuation (RMSF), Solvent Accessible Surface Area (SASA), and Radius of Gyration (Rg).

### 2.4 Statistical analysis

The results were elicited with triplicate value and expressed as mean ± SD. All the data were analysed statistically by Student t-test and two-way ANOVA using GraphPad Prism 5. A p-value <0.05 was considered statistically significant.

## 3.0 Results

### Cytotoxic effects of TMZ and FU on glioma cells

In order to validate the cytotoxic effect of the drugs, both individually and in combination on glioma cells, MTT assay was performed wherein, the cells were treated individually with varying concentrations of the drugs namely 10, 50, 100, 150, 200, 250, and 300 µM. In the present study, both the chosen drugs (TMZ and FU) had depicted a significant anti-proliferative effect against glioma cells. Among the two drugs employed for the current study, individually, TMZ had displayed a prominent growth inhibitory effect in a stable dose-dependent fashion (IC_50_: 50 ± 0.003 µM). However, FU as an individual drug showed only a moderate cytotoxic effect as evident from its IC_50_ value (150 µM± 0.02 µM) (Fig 2A).

Having determined the individual IC_50_ values of the drugs, we then assessed the effect of the combinatorial treatment on glioma cells. For this, the IC_20_ of TMZ (identified to be more potent based on individual IC_50_ values) was kept constant and the concentration of FU was varied. After 24 hours of treatment, we observed that the exposure to combinatorial treatment had sensitized glioma cells more significantly (P<0.05), than the effect observed as mono-therapeutic regimes. To note, when administered as a combinatorial treatment, the concentration of TMZ and FU was reduced to 1.2 µM and 89.04 µM respectively (Fig 1A). This data highlights the fact that FU can elevate the sensitivity of the cells to TMZ, at a minimal dosage. After 24 hours of treatment, the IC_50_ values of TMZ-alone, FU-alone and TMZ+FU in C6 glioma cells were found to be 50 ± 0.003 µM,150 ± 0.02 µM and 89.04 ± 0.05 µM respectively (Fig 1A).

**Fig 1:**
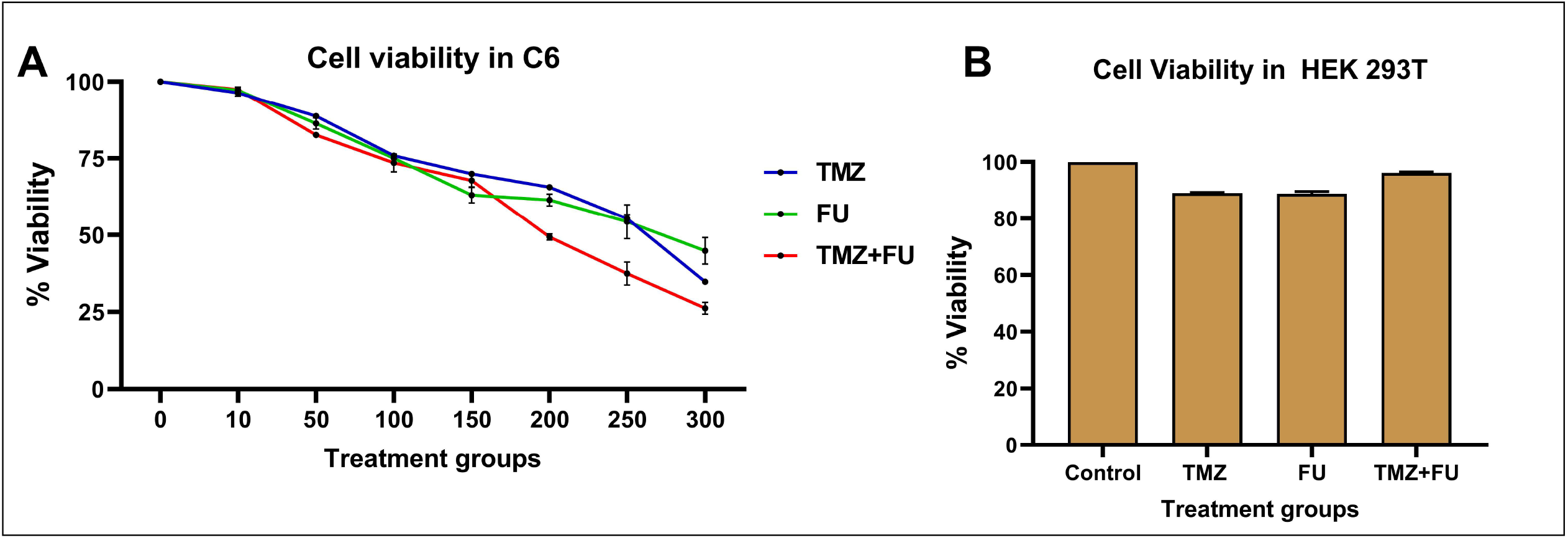
Cytotoxicity analysis of TMZ and FU, individually and in combination on (A) C6 and (B) HEK293T cells. Data points represent mean ± SD from triplicate.

**Fig 2:**
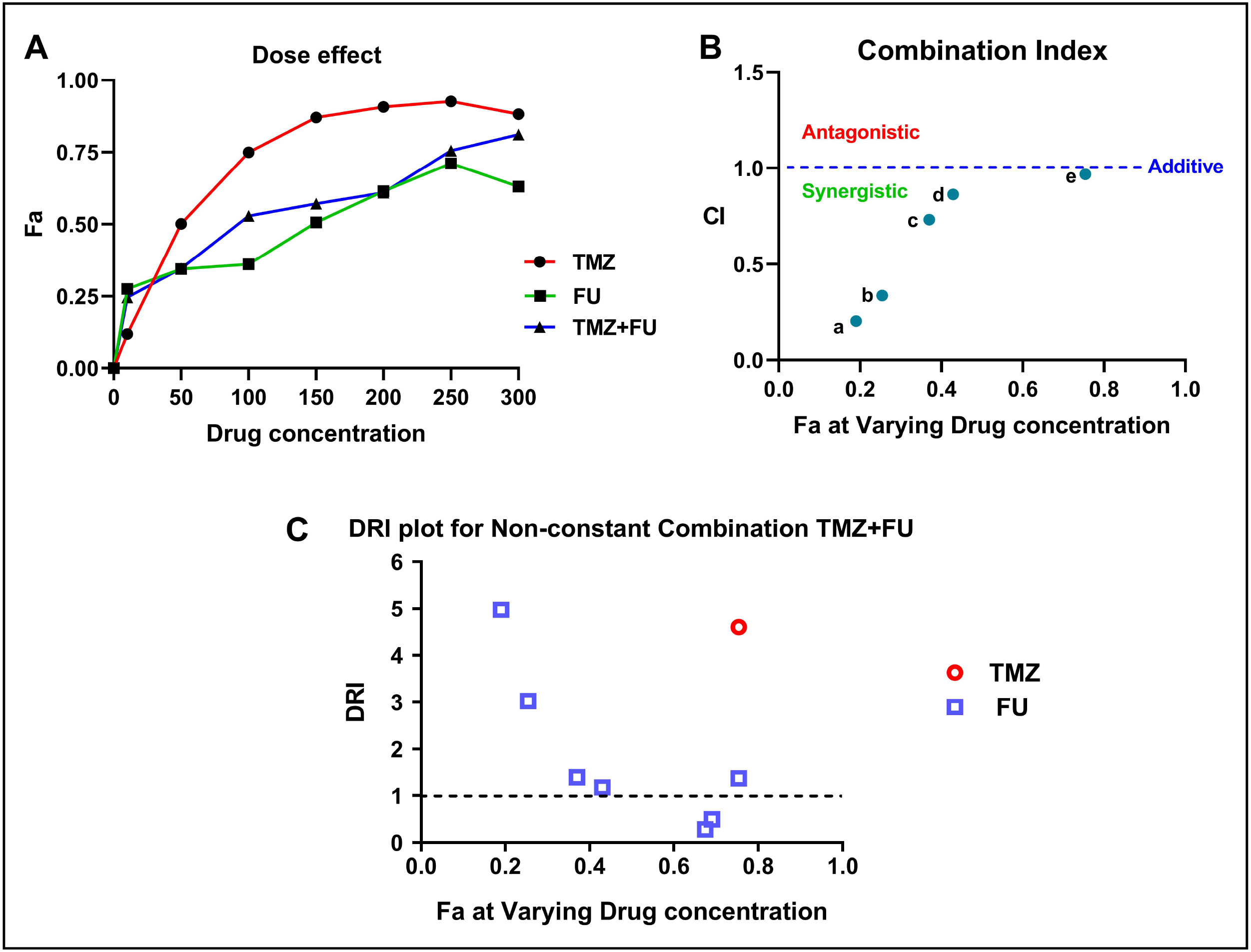
Combination Index Analysis using Compusyn: (A) Dose-effect curves for TMZ, FU and TMZ+FU after 24 hours of treatment. (B) Combination index plot: The combination index is plotted as a function of Fa. (C) Dose reduction index plot for combination: Dose reduction index values at different Fa values for each drug in the combination.

Further, in order to determine whether the individual and dual-drug treatment of TMZ and FU were toxic to normal cells, cytotoxicity was evaluated on HEK293T, a normal transformed cell line. We observed that the IC_50_ values obtained for individual and dual-drug treatment (TMZ+FU) on HEK293T were higher than that of glioma cells. These findings suggest that individual and combinatorial treatment of TMZ+FU was found to be more cytotoxic and sensitive to glioma cells, rather than the normal cells (Fig 2B).

### Selectivity index (SI)

Selectivity index (SI) of TMZ, FU and their combination of TMZ+FU were evaluated from the cytotoxicity assay performed on C6 and HEK293T cells. Drugs possessing higher SI values are considered to be more efficacious against cancer cell lines at concentrations obtained below the cytotoxic concentration of normal cells. The SI values from the present study revealed that the IC_50_ of TMZ in HEK293T cells was 5-fold greater than in C6 and IC_50_ of FU in HEK293T was 2-fold greater than in C6 cells, whereas combination of TMZ and FU showed a fold change of 2.24 greater in HEK293T, than that of C6 cells (Table 3).

**Table 3:** Selectivity index (SI) analysis

| Drugs used | IC<sub>50</sub> value in C6 glioma (μM) | IC<sub>50</sub> value in HEK293T cells ( μM) | Selectivity Index (SI) |
| --- | --- | --- | --- |
| TMZ | 50 | 250 | 5 |
| FU | 150 | 300 | 2 |
| TMZ +FU | 89.04 | 200 | 2.24 |

### Synergistic effects of TMZ and FU on glioma cell proliferation

To determine whether the cytotoxic effects of the dual-drug combination (TMZ+FU) were additive, synergistic or antagonistic, normalized isobologram and statistical combination index (SCI) for non-constant ratio combination design was assessed. CI values were calculated by the Compusyn software according to the recommendations of Chou-Talalay, wherein combinatorial-drug treatment (TMZ+FU) exhibited strong synergistic effect at an IC_50_ value of 89.04 µM (Fig 1A and 2A), CI index of 0.62 (Fig 2B). Besides, DRI value of the dual-drug combination was obtained more than 1, at its IC_50_ value, suggesting a favourable synergism (Fig 2C). These results suggest that the combinatorial-drug treatment of TMZ and FU, exhibited maximum synergistic cytotoxicity against glioma cells.

One of the major hallmarks for glioma pathogenesis is migration, which is a key contributing factor for most of the treatment failures. Inhibition of glioma proliferation of the combinatorial treatment drove us to investigate if this dual-drug cocktail could also hamper glioma migration. Thus, we performed wound healing assay and measured the cell-free area at 0 and 24 hours, following exposure to the drugs. Though the efficacy of TMZ (Fig 3B) and FU (Fig 3C) as individual drugs was relatively modest on sustaining the cell-free area, the combinatorial treatment (TMZ+FU) had rarely promoted the migration of glioma cells, with the cell-free area of micro-metre (µM). This highlights the fact that the combinatorial treatment (TMZ+FU) besides exhibiting significant anti-proliferative effect, can also inhibit the mobility of glioma cells (Fig 3D).

**Fig 3:**
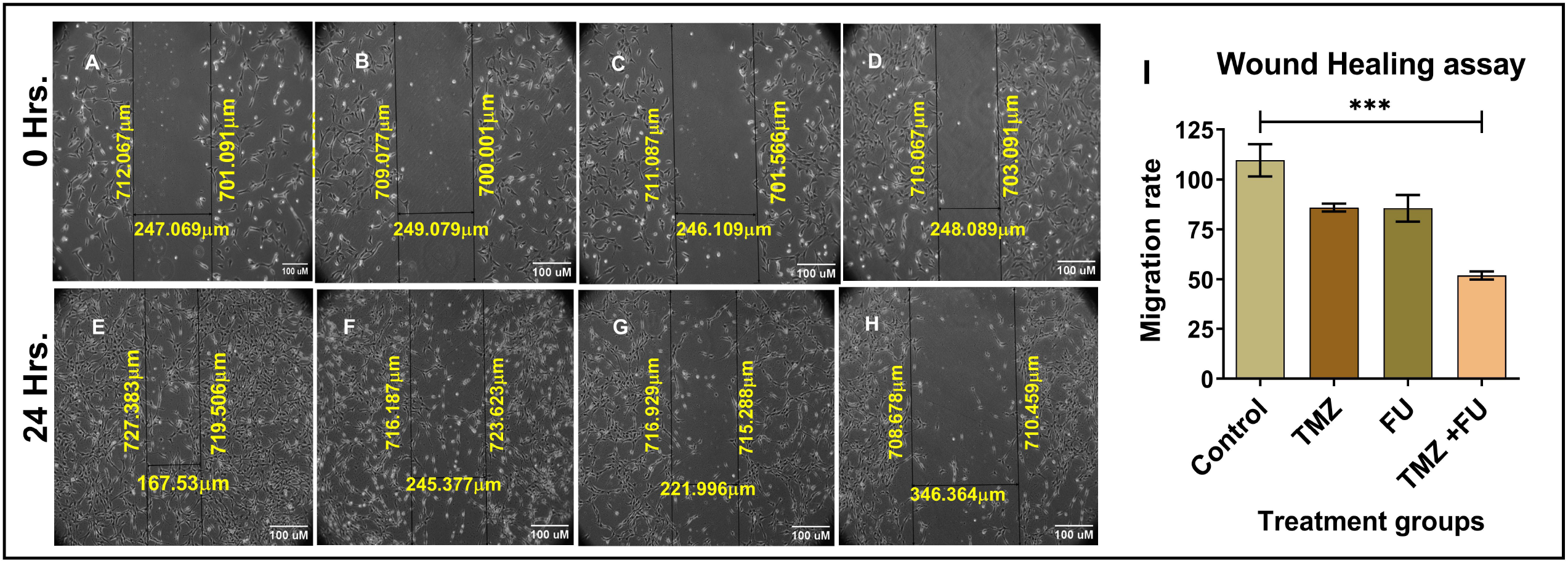
Inhibition of cell migration under the different treatment groups- (A & E) Control, (B &F) TMZ, (C & G) FU and (D & H) TMZ + FU, for 0hrs and 24 hrs, respectively. (I) Percentage of cell migration was calculated on the width of remaining wound surface area relative to the initial wound surface area. *** indicates p≤0.001, compared with control.

### Combinatorial Effect of TMZ and FU on apoptosis on C6 glioma cells

With the aim of understanding the inhibitory mechanism elicited by TMZ, FU or their combination on glioma cells, we analyzed the induction of apoptosis at morphological level by few staining procedures. This is due to the fact that, evasion of apoptosis is a distinct feature of most of the cancer cells including glioma.

Firstly, we performed H&E staining, wherein we found that treatment with TMZ-alone had endowed a moderate number of apoptotic cells (49.6%, Fig 4A) with shrinking nuclei and the cells had very poor demarcation of the nucleus and cytoplasm, when compared with that of control (untreated) cells. FU treatment on the other hand, resulted only a few apoptotic cells (Fig 4B), while most of the cells had a clear cytoplasm and nucleus, denoting that FU as an individual drug had moderate apoptosis induction potency but was not as effective as the standard drug, TMZ (33.06%). Interestingly, cells which were exposed to the combinatorial treatment (TMZ+FU) portrayed a large number of apoptotic cells (84.11%) with cell shrinkage (Fig 4D), and their viability was drastically reduced. Also, the cells had displayed marginated, condensed and aggregated nuclei, which all are the morphological characteristics of apoptosis.

**Fig 4:**
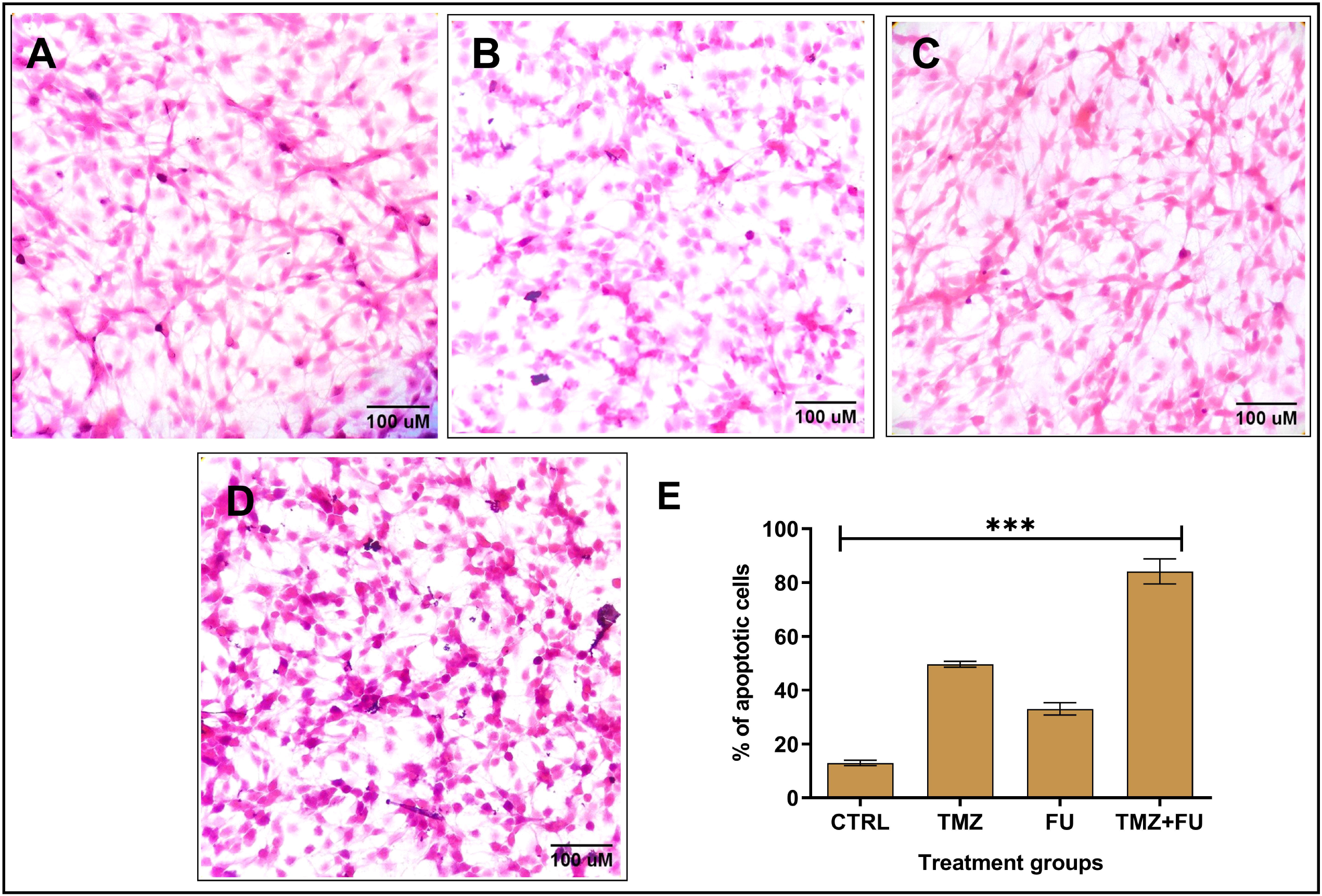
Apoptotic cell death observed using Hematoxylin and Eosin (H &E) stain under different treatment groups on C6 glioma cells (A) Control, (B) TMZ, (C) FU, (D) TMZ+FU and (E) Percentage of cells was calculated as total cellular shrinkage and loss of integrated density. *** indicates p<0.001, compared with control.

Following affirmation that the combinatorial treatment restrained glioma growth and induced apoptosis by H&E, we were also interested to distinguish the cells at different stages of apoptosis following administration of these drugs. For this purpose, AO/EtBr stain was employed. AO penetrates the live cells or cells in the early stage of apoptosis and stains them yellow-green, while EtBr is only permeable to the late-apoptotic cells with compromised cell membrane, emitting orange-red fluorescence. In accordance with this principle, our staining results revealed that in individually TMZ-treated cells (Supplementary Fig1B), a large number of cells at the early apoptotic stage (24%), and emitting yellow-green fluorescence were observed. On the other hand, treatment with FU (Supplementary Fig 1C) also resulted in a large number of early apoptotic cells and few late apoptotic cells (32.3%), signifying the early apoptotic effect of FU. Interestingly, combined administration of these drugs as a dual-drug treatment resulted in a large number of late-apoptotic cells (45%), emitting orange-red fluorescence (Supplementary Fig 1D). Also, a small proportion of the cells displayed an uneven orange-red fluorescence pattern, with increased cell volume, thereby denoting that the cells had undergone necrosis.

Following the analysis of cytoplasmic changes induced by the combinatorial treatment via H&E and AO/EtBr stains, we then employed the nuclear stain, DAPI to evaluate the alterations induced by TMZ, FU and their combination at nuclear level. Following staining, cells assigned as control, depicted a clear and distinct nucleus. As expected, the nuclear morphology in the treated cells was altered in all the treatment groups employed for our study. Among the individual drugs, treatment with TMZ endowed 70.48 % of apoptotic cells with condensed and marginated nuclei (Supplementary Fig 2B). However, treatment with FU resulted only in 67.28% of apoptotic cells (Supplementary Fig 2C) with condensed nuclei. The most significant nuclear alterations were apparent in the combinatorial treatment (TMZ+FU), wherein 73.3% of apoptotic cells with highly shrinked nuclei and patterns of nuclear fragmentation were observed (Supplementary Fig 2D). Taken together, our staining results suggested that the combinatorial treatment was bestowed with the potency to induce apoptosis at both cytoplasmic and nuclear level, besides suppressing glioma proliferation and migration.

### Dual-drug combination modulated the gene expression levels of inflammatory and pro-apoptotic markers in C6 glioma cells

Observations from the staining techniques led us to investigate the underlying molecular mechanism of this combinatorial treatment on glioma cells. Hence, we analysed the relative mRNA expression levels of inflammatory and pro-apoptotic genes namely IL-6, TLR-4, MYD88, IFN-γ, STAT3, NOS-2 and caspase-3 by qRT-PCR analysis. It is worth noting that the inflammatory genes IL-6, TLR-4, MYD88, NOS2, STAT3 and IFN-γ were aberrantly expressed in glioma’s and may contribute to tumor progression and chemo-resistance. Treatment with TMZ and FU, as individual drugs, had significantly down-regulated the expression levels of the inflammatory genes while concomitantly increasing the mRNA expression of caspase-3 to a considerable extent. Interestingly, among all of the experimental groups, the combinatorial treatment of TMZ+FU displayed a significant decrease in the fold change of inflammatory genes while up-regulating the fold change of caspase-3 (Fig 5), thereby denoting that this combination elicited apoptosis by down-regulating the inflammatory signaling cascade.

**Fig 5:**
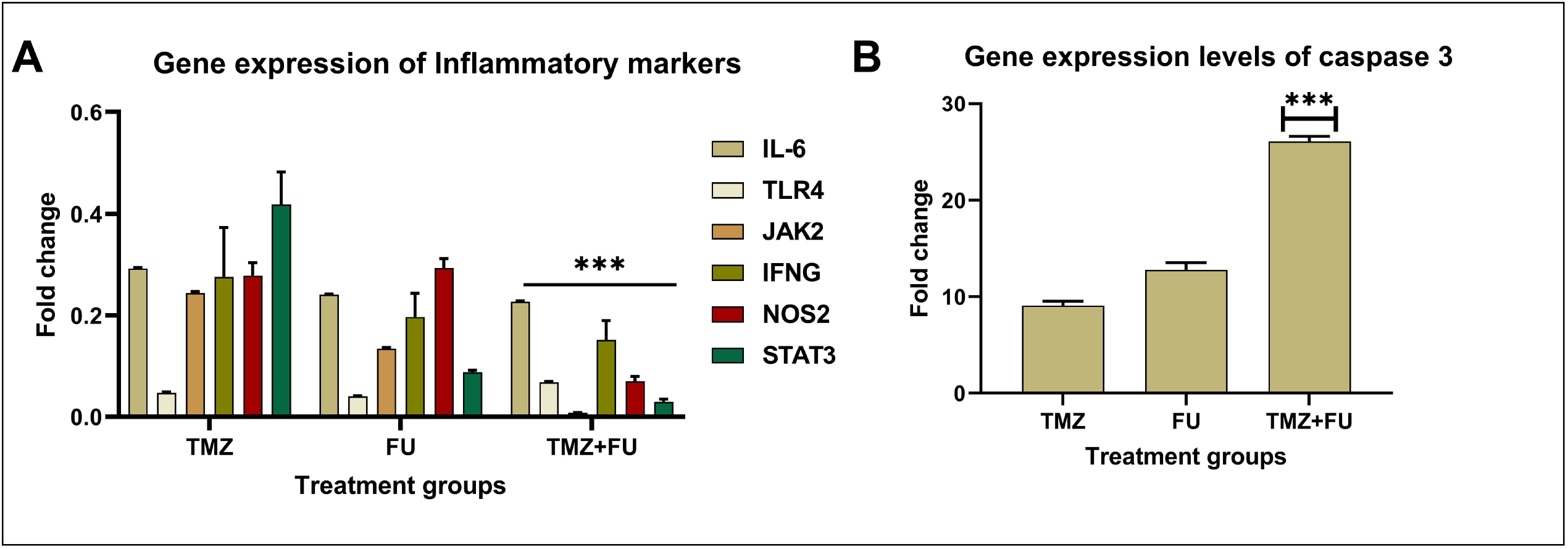
Anti-glioma effects of FU, both alone and in combination with TMZ on the mean fold change (Linear scale) of inflammatory genes (A) IL-6, TLR4, MYD88, IFN-γ, NOS2, STAT3 and apoptotic gene (B) caspase-3 in C6 glioma cells. *** indicates p<0.001 in comparison to untreated control.

### Molecular docking

#### Active site prediction

The active sites were predicted using Sitemap prediction tool (Schrodinger) and have been listed in the Supplementary Table 1.

#### Molecular Docking analysis

In order to predict the interaction of the ligands with the key drivers of inflammation in glioma, docking analysis was performed for the following targets namely, IL-6, TLR4, JAK2, STAT3, IFN-γ and NOS2. The detailed interaction profile of the ligands with inflammatory proteins has been tabulated in Table 5.

Interestingly, we found that, all of the anti-tumor ligands viz., TMZ, MTIC, AIC and FU interacted with IL-6 by H-bonds predominantly. With regard to the binding energies, we observed that, TMZ, MTIC and AIC formed more number of H-bonds and exerted a binding energy of -23.953, -13.480 and -28.148 Kcal/mole (Table 4). Although FU exhibited only two H-bonds with IL-6 (Fig 8), it depicted a highest binding energy of -28.902 Kcal/mole (Table 4). This could be due to the presence of other non-bonding interactions, which could have facilitated the stronger interaction of FU with IL-6 (Fig 6).

**Table 4:** Interaction profile of all the inflammatory targets against the four ligands

| Ligands | Interaction type | Functional moieties<br>{ } - Interactions with same amino acid residues | Amino acid residues | Binding Energy (Kcal/mole) |
| --- | --- | --- | --- | --- |
| **IL-6** |  |  |  |  |
| **TMZ** | H-bonds (4) /salt bridge (1) | NH<sub>2</sub><sup>+</sup> (at 1<sup>st</sup> position of tetrazine ring), NH (6<sup>th</sup> position of tetrazine ring), O<sup>-</sup> (7<sup>th</sup> position) of carboxamide) and NH<sub>3</sub><sup>+</sup> (8<sup>th</sup> position) of carboxamide in imidazole ring/ | Leu90, Leu89, Ile87, Ser186/ Lys91 | -23.953 |
| **MTIC** | H-bonds (3) | NH<sub>2</sub><sup>+</sup> (1<sup>st</sup> position), N<sup>+</sup>H<sub>2</sub> (3<sup>rd</sup> position), O<sup>-</sup> (4<sup>th</sup> position) of carboxamide and NH<sub>2</sub> (5<sup>th</sup> position) of tetrazine ring | Lys91, Leu90, Arg74 | -13.480 |
| **AIC** | H-bonds (4) | OH (at 1<sup>st</sup> position), NH<sub>2</sub><sup>+</sup> (2<sup>nd</sup> position), {NH<sub>2</sub><sup>+</sup> (4<sup>th</sup> position), substituted chain (at 5<sup>th</sup> position, NH-3<sup>rd</sup> position)} of imidazole ring | Gly77, Cy78 and Gln85, | -28.148 |
| **FU** | H-bonds (2) | OH (4-Hydroxy), O<sup>-</sup>SO<sub>3</sub> (at 5<sup>th</sup> position) (sulphate) of Fucose ring / | Leu89, Leu90 | -28.902 |
| **TLR4** |  |  |  |  |
| **TMZ** | H-bonds (3)/ salt bridge (3) | {NH<sup>+</sup> at 4<sup>th</sup> position and N<sup>+</sup>H<sub>2</sub> at 6<sup>th</sup> position, N<sup>+</sup>H<sub>3</sub> at 8<sup>th</sup> position} of imidazole ring/ {NH<sup>+</sup> at 4<sup>th</sup> position and {N<sup>+</sup>H<sub>2</sub> at 6<sup>th</sup> position}, N<sup>+</sup>H<sub>3</sub> at 8<sup>th</sup> position} } of imidazole ring | Asp 208, Glu229/ Asp180, Asp208 and Glu229 | -27.448 |
| **MTIC** | H-bonds (3) /salt bridge (2) | NH<sub>2</sub><sup>+</sup> (at 1<sup>st</sup> position), {NH<sub>2</sub><sup>+</sup> (at 3<sup>rd</sup> position), NH<sub>3</sub><sup>+</sup> (at 4<sup>th</sup> position-of imidazolidin)}/ NH<sub>2</sub><sup>+</sup> (at 1<sup>st</sup> position), NH<sub>2</sub><sup>+</sup> (at 3<sup>rd</sup> position) of imidazole ring | Glu229, Asp208/ Glu229 and Asp208 | -40.175 |
| **AIC** | H-bonds (5) /salt bridge (3) | N<sup>+</sup>H<sub>3</sub> (1<sup>st</sup>), NH<sub>2</sub><sup>+</sup> (2<sup>nd</sup>), NH<sub>2</sub><sup>+</sup> (4<sup>th</sup>), 5<sup>th</sup>-substituted chain (1<sup>st</sup> and 3<sup>rd</sup> NH-NH) of imidazole ring/ NH<sub>2</sub><sup>+</sup> (2<sup>nd</sup>) of imidazole ring | Glu229, Asp208 / Glu229, Asp180, Asp 208 | -25.524 |
| **FU** | H-bonds (2) /salt bridge (1) | O-SO<sub>3</sub><sup>-</sup> (at 3<sup>rd</sup> position) and OH (5-hydroxy) of Fucose ring | Arg233, Arg337/ Lys263 | -23.074 |

| **JAK2** |  |  |  |  |
| --- | --- | --- | --- | --- |
| **TMZ** | H-bonds (3)/<br>salt bridge<br>(2) | O<sup>-</sup>(4<sup>th</sup> position) of carboxamide, NH<sub>2</sub><sup>+</sup> at 7<sup>th</sup> and 8<sup>th</sup> position of imidazole ring – (OH & NH<sub>2</sub>)/ O<sup>-</sup> (4<sup>th</sup> position) of carboxamide and NH<sub>2</sub><sup>+</sup> at 7<sup>th</sup> position of imidazole ring | Leu824, Cys905, Asp908, Asp908/ Arg912 and Asp908 | -32.530 |
| **MTIC** | H-bonds (7) | N<sup>+</sup>H<sub>2</sub> (at 1<sup>st</sup> position), N<sup>+</sup>H<sub>2</sub> (3<sup>rd</sup> position), OH & NH<sub>2</sub> (at 4<sup>th</sup> position) of imidazole ring | Tyr900, Ser 573, Leu583, His510, Tyr900 and Pro902 | -42.885 |
| **AIC** | H-bonds (7)/<br>salt bridge<br>(1) | NH<sub>2</sub> and OH (1<sup>st</sup> position), NH and NH<sub>2</sub><sup>+</sup> (2-(3-methyl-2λ<sup>4</sup>-triazenyl), NH (3<sup>rd</sup> position) and NH<sub>2</sub><sup>+</sup> (5<sup>th</sup> position) of imidazole ring/ NH<sub>2</sub><sup>+</sup> (5<sup>th</sup> position) of imidazole ring | Asp963, Arg949, Asn826, Leu824/ Asp963 | -33.159 |
| **FU** | H-bonds (4) | O-SO<sub>3</sub><sup>-</sup> (3<sup>rd</sup>) of sulphate, OH at 4<sup>th</sup> and 5<sup>th</sup> position of Fucose ring | His510, Arg912, Ser903, Pro902 | -29.593 |
| **STAT3** |  |  |  |  |
| **TMZ** | H-bonds (3)/<br>salt bridge<br>(1) | NH (at 1<sup>st</sup> position), NH<sub>3</sub><sup>+</sup> (at 9<sup>th</sup> position), of imidazole ring/ O<sup>-</sup> at 3<sup>rd</sup> position (oxidanyl group) of imidazole ring | Glu455, His457/ Lys244 | -17.346 |
| **MTIC** | H-bonds (4)/<br>pi (π)-cation<br>(1) | {NH<sub>2</sub> (at 1<sup>st</sup> position), NH<sub>2</sub>} (2<sup>nd</sup> position at imidazolidin-4-yl), NH (5<sup>th</sup> position)/ NH<sub>2</sub><sup>+</sup> (at 3<sup>rd</sup> position) of imidazole ring | Gln247, Val323 and His457/His457 | -28.236 |
| **AIC** | H-bonds (4)/<br>salt bridge<br>(2)/ pi (π)-<br>cation (1) | NH<sub>3</sub><sup>+</sup> (at 1<sup>st</sup> position), {NH<sub>2</sub><sup>+</sup> (4<sup>th</sup> position), substituted chain (5<sup>th</sup> position of 1<sup>st</sup> and 3<sup>rd</sup> NH-NH)} of imidazole ring /NH<sub>2</sub><sup>+</sup> at 2<sup>nd</sup> position of imidazole ring/ NH<sub>2</sub><sup>+</sup>(2<sup>nd</sup> position of substituted chain) | Glu455, Asp371/ Glu455, Asp371/ His457 | -20.667 |
| **FU** | H-bonds (2) | OH at 3<sup>rd</sup> position and O-SO<sub>3</sub><sup>-</sup> (at 5<sup>th</sup> position) of Fucose ring | Thr456, His457 | -23.658 |
| **IFN-γ** |  |  |  |  |
| **TMZ** | H-bonds (4) | NH (at 7<sup>th</sup> position), OH & NH<sub>2</sub> (at 9<sup>th</sup> position) of imidazole ring | Cys178, Glu175, Thr172 | -31.600 |
| **MTIC** | H-bonds (3) | {NH (4<sup>th</sup> position), (carboxamide), NH<sub>3</sub><sup>+</sup> (5<sup>th</sup> position)}, of imidazole ring | Cys178, Thr172 and Thr172 | -24.403 |
| **AIC** | H-bonds (7)/<br>salt bridge<br>(1) | NH (at 1<sup>st</sup> position), NH-NH (at 1<sup>st</sup> and 3<sup>rd</sup> position), NH<sub>3</sub><sup>+</sup>(3<sup>rd</sup> position), NH<sub>2</sub><sup>+</sup>(4<sup>th</sup> position) of imidazole ring/ NH<sub>2</sub><sup>+</sup>(4<sup>th</sup> position) of imidazole ring | Cys178, Asp176, Asp176, Glu175 and Gln173 | -36.282 |
| **FU** | H-bonds (3) | OH (3,4-dihydroxy), O=SO<sub>3</sub><sup>-</sup> of Fucose ring | Phe134, Ile152 | -23.932 |

| **NOS2** |  |  |  |  |
| --- | --- | --- | --- | --- |
| **TMZ** | H-bonds (2)/<br>pi ($\pi$)-cation<br>(2) | {NH<sub>2</sub> and OH} (at 8<sup>th</sup> position)/NH<sub>2</sub><sup>+</sup> at 7<sup>th</sup><br>position of imidazole ring | Asn364, Phe363 and<br>Trp188 | -24.211 |
| **MTIC** | H-bond (1)/<br>pi ($\pi$)-cation<br>(2) | NH<sub>2</sub> (at 2<sup>nd</sup> position)/ NH<sub>2</sub><sup>+</sup> (4<sup>th</sup> position) and C-at<br>5<sup>th</sup> position of imidazole ring | Asn364/ Trp188 and<br>Phe363 | -20.902 |
| **AIC** | H-bonds (6)/<br>salt bridge<br>(1) | {NH<sub>2</sub> (at 1<sup>st</sup> position), (1<sup>st</sup> and 3<sup>rd</sup> position NH-NH at 2<sup>nd</sup> position of substituted chain,), {N<sup>+</sup>H<sub>2</sub><br>(3<sup>rd</sup> position), N<sup>+</sup>H<sub>2</sub> (5<sup>th</sup> position) and OH (1<sup>st</sup>)} of<br>imidazole ring/ N<sup>+</sup>H<sub>2</sub> (3<sup>rd</sup> position) of imidazole<br>ring | Trp366, Glu361/<br>Glu361 | -33.902 |
| **FU** | H-bond (1) | O<sup>-</sup>-SO<sub>3</sub> bonded at 2<sup>nd</sup> position of Fucose ring | Tyr485 | -24.978 |

**Fig 6:**
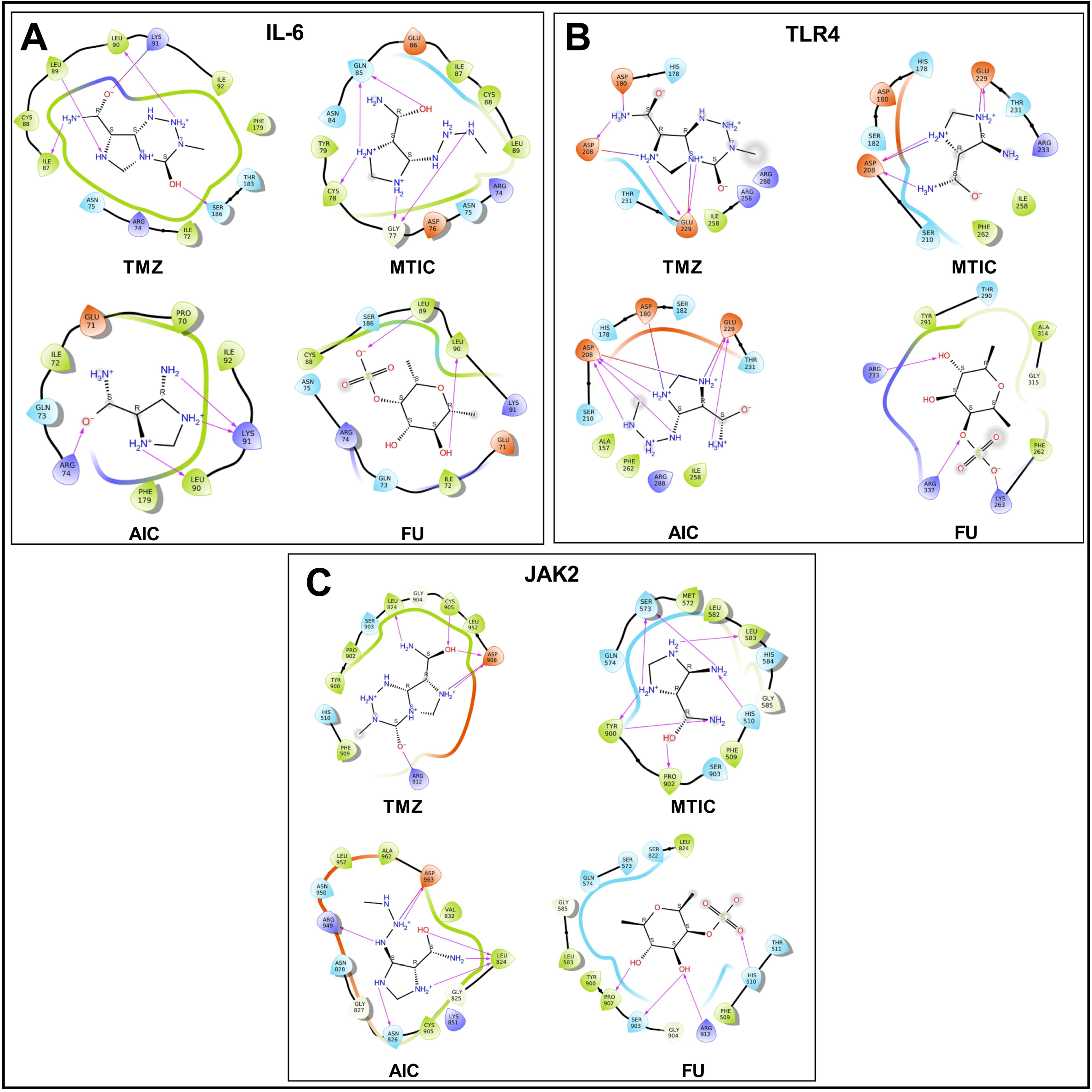
Interaction profile of (A) IL-6, (B) TLR4, (C) JAK2 with ligands-TMZ, MTIC, AIC and FU

In the case of TLR4, we observed that TMZ, MTIC, AIC and FU interacted through H-bonds and salt bridge formation (Fig 6), with a binding energy of -27.448, -40.175, -25.524, and -23.074 Kcal/mole, respectively (Table 4). Among the anti-tumor ligands, MTIC, despite possessing lesser number of H-bonds and salt bridges, depicted the highest binding energy (Fig 6).

A similar pattern of interaction was observed in the case of JAK2, whereby, MTIC exhibited the highest binding energy of -42.885 Kcal/mole, while the remaining ligands-TMZ, AIC and FU exerted a binding energy of -32.530, -33.159 and -29.593 Kcal/mole, respectively (Table 4).

Likewise, the docking analysis for STAT3 revealed that, MTIC depicted the highest binding energy of -28.236 Kcal/mole, while TMZ, AIC and FU exhibited binding energy of, -20.667, 17.346 and -23.658 Kcal/mole respectively (Fig 7, Table 4). The binding energy of MTIC could be due to the formation of π-cationic interaction along with H-bond that contributed to the overall stability, when compared with the other anti-tumor ligands (Fig 7).

**Fig 7:**
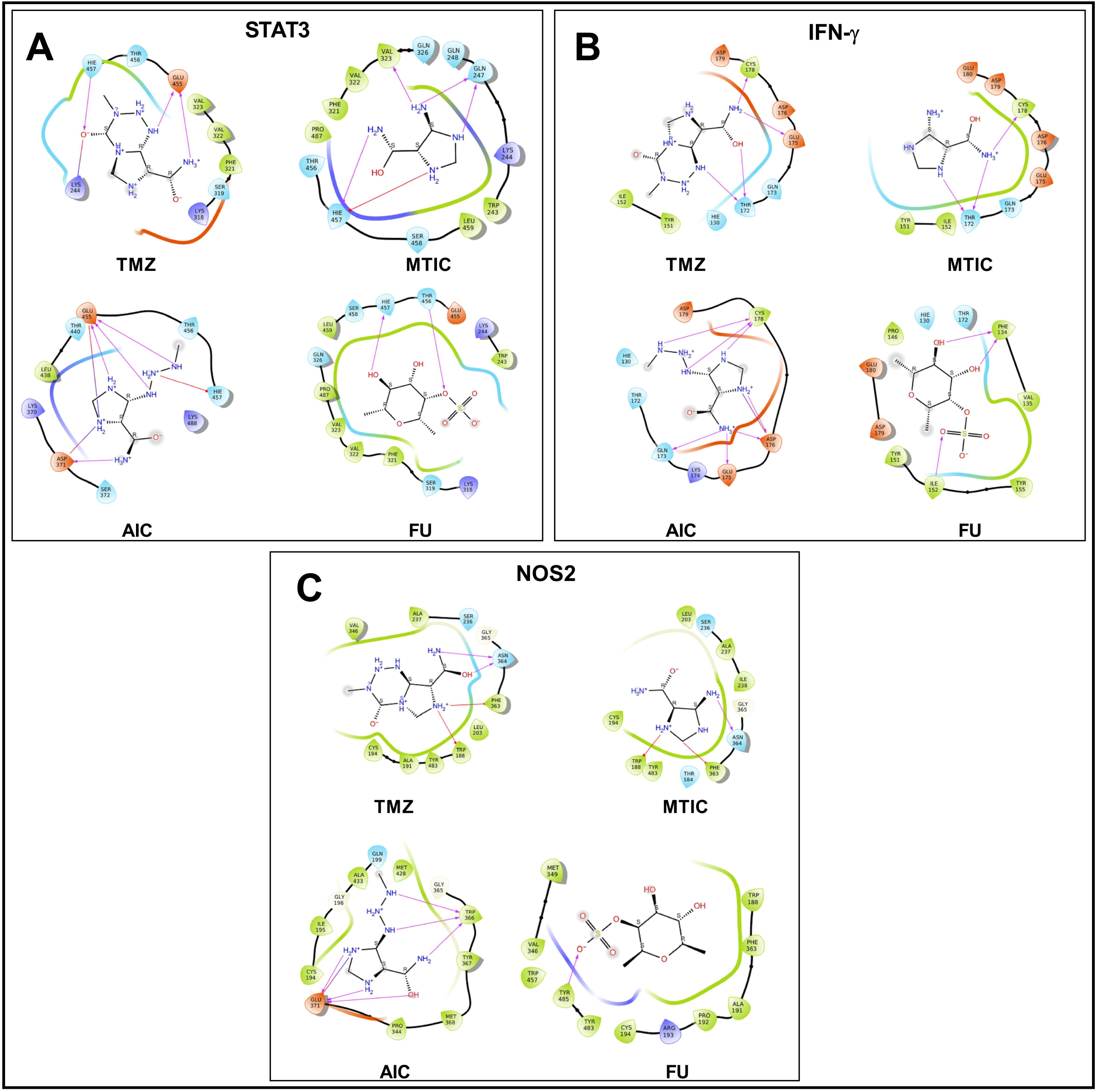
Interaction profile of (A) STAT3, (B) IFN-γ and (C) NOS2 with ligands TMZ, MTIC, AIC and FU.

In the case of IFN-γ, docking results revealed that AIC depicted the maximum interaction profile with a binding energy of -36.282 Kcal/mole (Fig 7), while TMZ, MTIC and FU exerted a binding energy of -31.600, -24.403 and -23.932 Kcal/mole respectively (Fig 7, and Table 4).

With the most crucial target, NOS2, docking analysis revealed that AIC exhibited a highest binding energy of -33.902 Kcal/mole (Fig 7), following, FU, TMZ and MTIC of -24.978, -24.211 and -20.902 Kcal/mole (Fig 7). This accounts for the presence of maximal number of H-bonds and salt bridge formation, as compared to other ligands and hence the maximum stability (Table 4).

To summarize, the complexity of bonding between the ligand-target protein complexes denotes the strength of molecular docking. From our visual analysis, based on the binding energy and docking score, it was evident that TMZ and its metabolites, MTIC and AIC were the best ligands to form complexes with TLR4, JAK2, STAT3, IFN-γ and NOS2, while FU exhibited highest interaction with IL-6. However, further investigations were warranted for verification of the stability of these ligand-protein complexes obtained by docking analysis.

#### Molecular Dynamics Simulation (MDS)

Following identification of the docked complexes and affirmation of their interaction profile, we then determined the stability of these interactions using Molecular Dynamics Simulation (MDS). MDS was evaluated in terms of RMSD, RMSF, Rg and SASA against each of the docking complexes.

Among the MDS parameters, RMSD is used to measure the average change in displacement of atoms of a particular frame with respect to a reference frame. In general, the target reaches stability in few pico-seconds (ps). Initially RMSD was performed for IL-6 whereby we found that, TMZ, MTIC and AIC depicted a stable interaction (0.5-0.59 nm) with the target protein till 20000 ps, while FU exhibited a slight increase in the RMSD value (0.66 nm) from 15000 ps (Fig 10). Similar to their interaction with IL-6, the ligands TMZ, MTIC and AIC showed a rise from (0.6-0.9 nm), following consistent RMSD plot (0.9 nm) for TLR4 as well, while FU portrayed a mild increase in RMSD value (1.2 nm) from 10000 ps. Interestingly, all of the four ligands displayed an overall stability for JAK2 and STAT3 at 20000 ps (0.4 nm). On the other hand, RMSD plots of IFN-γ and NOS2 revealed a stable RMSD value (0.2 nm) for all the chosen ligands, except AIC, which showed a elevation in the RMSD plot (IFN-γ - 0.8 nm at 15000 ps; NOS2-0.6 nm at 10000 ps) (Supplementary Fig 3). Based on the RMSD studies, we concluded that all of the chosen protein-ligand complexes had attained structural stability against all of these docked complexes. To further verify the ligand-protein stability, we then evaluated the RMSF value for the ligand-protein interactions. RMSF plot enables to predict if the ligand-target interaction is sufficiently stable and provides basis for the flexibility of the conformation. RMSF fluctuation is often connected to RMSD plot, wherein more RMSF fluctuation yields less stable RMSD plot. Interestingly, we found that among the ligands that interacted with IL-6, TMZ, MTIC and AIC exhibited the minimum fluctuation (<0.4 nm) while FU depicted the maximum fluctuation (0.7 nm) (Fig 10). In the case of TLR4, RMSF plot revealed that among all of the four ligands interacted, TMZ, AIC and FU showed the minimum fluctuation (<0.8 nm) while MTIC exhibited maximum fluctuation of 1.0 nm. Conversely, the RMSF plots of JAK2 revealed minimal fluctuation in TMZ (0.4 nm) and maximum fluctuation for MTIC, AIC and FU (0.8 nm). On the other hand, amongst all of the docked complexes, MTIC, AIC and FU depicted a minimal fluctuation (<0.4 nm) with IFN-γ, while TMZ demonstrated a slight increase in RMSF (at 200 residues). With regard to their interaction with NOS2, TMZ, MTIC and FU exhibited a minimum fluctuation of <0.5 nm, while AIC showed the maximum fluctuation (2.5 nm). Taken together, in consensus with our RMSD results, the RMSF affirmed that the ligand-protein complexes were sufficiently stable and had low flexible conformation (Supplementary Fig 3).

SASA is used to determine the surface area on a biomolecule that is accessible to solvent molecules. SASA plot denotes the ligand-protein interaction in the solvent. In general, low SASA value denotes that the target protein is less available for interaction with the solvent molecules like water; High SASA value on the other hand, implies that the target protein is widely accessible by the solvent molecules. Interestingly, the SASA plots of the interactions between all the four ligands and IL-6 revealed a low SASA value (100-140 nm^2^). Likewise, interaction between the ligands and TLR4 depicted a SASA value of 240-299 nm^2^. On the other hand, interaction of the anti-tumor ligands with JAK2, STAT3 and NOS2 exhibited a SASA value of 285-305 nm^2^, 285-305 nm^2^ and 205-225 nm^2^, respectively. In the case of IFN-γ, MTIC, AIC and FU exhibited a low SASA value of 100 nm^2^, while TMZ exhibited slightly higher SASA value of 220 nm. Taken together, our SASA analysis revealed that most of the chosen proteins depicted a low SASA score and had no folding effect on complex stability (Supplementary Fig 3).

Following elucidation of complex stability and surface exposure, the intrinsic dynamics of ligand-protein complexes were calculated by performing Rg on the docked complexes. It is well known that an increase in Rg values implies a decrease in the protein compactness leading to increased flexibility and less stability. Conversely, low Rg values on the other hand, denote stability and protein compactness. The Rg plots of all the chosen ligand-protein complexes are depicted in Fig 10. It can be seen that the Rg value of all the chosen ligands with IL-6 was measured between 1.55-1.70 nm with less fluctuation. In the case of TLR4, TMZ, MTIC and AIC depicted moderate high fluctuation and high Rg value between 3-3.4 nm at 10000 ps while FU depicted high fluctuation of 3.5 nm at 10000 ps. For JAK2, MTIC and AIC depicted low fluctuation with Rg value ranging from 2.7 to 2.74 nm, while TMZ and FU demonstrated a high fluctuation rate of 2.76-2.8 nm. However, all of these four ligands depicted a high fluctuation, while Rg value for STAT3 ranging from 3.42 to 3.54 nm. For IFN-γ, TMZ depicted the highest Rg value of 2.6 nm, while MTIC, AIC and FU showed Rg value below 2 nm with low fluctuation. Parallelly, for NOS2, all the four ligands exhibited a low fluctuation and Rg value from 2.3 to 2.36 nm. Thus, from our Rg scores, we concluded that the chosen ligand-protein complexes had depicted sufficient stability and compactness.

Overall, our *in silico* data strongly established the fact the chosen anti-tumor ligands had formed stable and compact docking complexes with the inflammatory proteins and may be probed further, to modulate the inflammatory pathway in experimental setup as an affirmation of our *in vitro* findings.

## 4.0 Discussion

Glioma, accounting for about 40% of the intra-cranial tumours, is often associated with a poor prognosis and dismal survival rate. The current cutting-edge treatment modality encompasses surgical resection of the tumor mass trailed by radio and chemotherapy employing TMZ, the gold-standard drug for glioma[14]. Though TMZ, as a mono-therapeutic agent had hailed a major breakthrough, its therapeutic efficacy has decreased over-years due to chemo-resistance. This puts forth a vital need for effective therapeutic strategies. Besides, several preclinical studies had showed that employing mono-therapeutic agents to target glioma were not efficacious and were not associated with a significant patient outcome [15]. This way, combination therapy has become the cornerstone of the current scenario, as it can bring about synergistic growth inhibitory effect while focusing on multiple gene targets at the same point. For instance, a combination of TMZ with thymoquinone, a natural product had showed synergistic inhibition of proliferation of glioma cells, by inducing apoptosis [16]. Pandey et al., reported that a cyclin-dependent kinase inhibitor, roscovitine, was known to potentiate the cytotoxicity of TMZ and reduced progression rate of glioma, either alone or in combination both *in vitro* and *in vivo* [17]. Above all, it has been put forward that, since TMZ is associated with considerable cytotoxic effects, synergizing TMZ with natural/ synthetic derivatives would bring about concurrent reduction in the dosage and would be a beneficial approach for glioma treatment.

In the search for potent drug candidates for combination therapy in glioma, derivatives of MNPs, which are less/non-toxic are often sought as they are associated with minimal side effects. FU, one such MNP, has been explored as a potential anti-tumor agent in several cancer cells. A recent study by Park et al, reported the plausible anti-inflammatory effects of FU through inhibition of NF-ƘB, MAPK and Akt activation in lipo-polysaccharide induced in BV2 microglial cells[18].

Inflammatory cytokines, are one of the major attributes in the pathogenesis of glioma, that contributes to tumor proliferation and sustenance, thereby promoting cell growth in glioma. Interleukin-6 (IL-6), a pro-inflammatory autocrine and paracrine cytokine possess tumor promoting and progressing effects in glioma[19]. It acts directly on tumor cells by binding to interleukin-6 receptor (IL-6R) or glycoprotein complex 130 (gp130 complex) and activates Janus Kinases (JAK) - STAT3. STAT3, which belongs to a family of transcriptional factors, is involved in proliferation, inflammation and tumorigenesis in most of the cancer types, including GBM. On the other hand, inflammation mediated by binding of toll-like receptor (TLR-4), can induce expression of nitric oxide synthase isoform 2 (NOS2) which leads to the production of nitric oxide. Owing to the importance of IL-6 mediated JAK2/STAT3 signalling pathway in tumorigenesis, therapeutic targeting of this pathway has elevated the role of stratified treatment approaches in most of the cancers, including glioma.

Previously, TMZ and FU were reported to exert potent anti-proliferative and growth inhibitory effects, individually, in various cancer cells. In accordance with those findings, in the present study, the dual-drug combination (TMZ+FU) was found to be non-toxic in normal cells, however, when this cocktail was administered on glioma cells, the drugs significantly hampered the proliferation of glioma than their individual counterparts (Fig 1 and 2). This may be attributed to the combination of the anti-proliferative effect of TMZ and growth inhibitory potency of FU. Further, to determine if such effective anti-proliferative effect was synergistic, we calculated the combination index for the dual-drug combination. We found that on average, 172.105-fold reduction of TMZ and 1.18-fold reduction of FU were sufficient to induce the median effect, while administering the drugs as a dual-drug combination. Moreover, the dual-drug combination along with individual drugs had a higher SI index, suggesting that the dual-drug combination selectively inhibited glioma cell growth (Table 3). Obtaining this synergistic anti-proliferative effect within such reduced drug dosage, would help to improve glioma prognosis and uphold patient survival. One interesting point to be noted in glial tumors is that, proliferation and migration are mutually exclusive. Alterations in the tumor micro-environment prompt glioma cells to “go/migrate” and re-settle to “grow/proliferate” in an adaptable environment[20]. This “Go and grow” phenomenon is of central importance in glioma, as it promotes tumorigenicity. Therefore, it is plausible that identification of chemo-therapeutic agents that inhibits the “Go” of glioma would help in halting the migration of tumor cells and subsequently inhibit glioma proliferation [21]. Earlier studies reported that individually TMZ had inhibited the migration of glioma *in vitro* in U251 [22]. Further, FU hampered proliferation and migration of liver cancer cell HepG2 [23]. In accordance with these findings, our results revealed that both the drugs had potent anti-migratory effect against glioma (Fig 3). Further, administration of these drugs as a combinatorial treatment had maintained the cell-free space to a significant level, thereby, halting the “Go” phenomenon and suppressing glioma proliferation (Fig 3). One possible mechanism for this anti-migratory effect exhibited by this dual-drug treatment would be due to suppression of the matrix metallo proteinases (MMPs), at least partially, by the combination of TMZ+FU. As an interesting note, the dual-drug combination of TMZ and FU regulated the expression of MMPs in BALB/C mice model (U87MG-fLuc transfected model) [24] and HT1080 fibro sarcoma cells [25]. However, profound molecular analysis is required to conclusively determine if the anti-migratory effect of this combinatorial treatment was due to the suppression of MMPs.

The anti-proliferative and the anti-migratory effects of the combinatorial drug treatment might be due to induction of apoptosis. In this perception, we perceived that our dual-drug combination (TMZ+FU) exhibited maximum alteration at nuclear and cytoplasmic level (Supplementary Fig 1 and 2), which can be triggered by activation of apoptotic signalling pathways, in comparison to control and individual-drug treated groups. Most cancer cells including GBM exhibits evasion of apoptosis and transforms to a high grade of malignancy that generates therapy-resistance [26]. Therefore, restoration of apoptotic pathway by therapeutic targets is one of the promising challenges faced in the field of anti-glioma drug development, while their exact mechanism of action has not been clearly demonstrated. It is well known fact that cell death by apoptosis, is mainly transmitted through cascades of caspase-mediated reactions. Caspases are prevalent chemotherapeutic or chemo preventive targets, from the perception of anti-cancer therapeutics [27]. Most of the anti-cancer agents derived from marine resources have shown to induce apoptosis by activation of caspase-3 [27].

Earlier studies reported that TMZ might induce apoptosis by mitochondrial translocation of STAT3, in a non-canonical dependent pathway [28]. In a similar study, it was reported that STAT3 suppression leads to enhanced sensitization of glioma by TMZ via down regulation of MGMT expression [29]. On the other hand, FU was also known to induce apoptosis by JAK/STAT3 signalling and promoting autophagy as well [30]. Also, FU was shown to induce apoptosis in liver cancer cell line HepG2 by down regulation of p-STAT3 [31]. In accordance with these findings, we observed that dual-drug combination of TMZ and FU led to enhancement of caspase-3 expression significantly, in comparison to individual drug treated and control groups. Thereof, we postulate that the combinatorial treatment of TMZ and FU augmented apoptosis by elevating the levels of caspase-3, possibly by down regulating the expression of JAK/STAT3 (Fig 5).

Further, to elucidate the possible binding mechanism of each of these ligands, TMZ, FU, MTIC and AIC, with the pro-inflammatory targets-IL-6, TLR4, JAK2, STAT3, IFN-γ and NOS2, individually, molecular docking was performed. Docking performed of TMZ, MTIC, AIC and FU at the active sites of the target proteins were analysed using parameters such as Glide based binding energy, hydrogen bond, π-cation, and salt bridge interactions. The binding energy revealed that FU-IL-6 exhibited highest binding energy, as compared to the other ligand-IL-6 complexes. Simulation studies revealed that each of the complexes (IL-6-TMZ, MTIC and AIC) were stable throughout the run time period of 20,000ps, which could be due to formation of H-bond (Supplementary Fig 3, Fig 6 and 7).

Further, we observed that (TLR4, JAK2 and STAT3)-MTIC; (IFN-γ and NOS2)-AIC binds strongly, as evident by their highest binding energy values. We found major л-cations interactions among STAT3-MTIC and AIC; NOS2-TMZ and MTIC complexes. Additionally, there were formation of salt bridges among (TLR4-TMZ, MTIC and AIC; JAK2-TMZ and AIC; STAT3-TMZ and AIC; IFN-γ-AIC; NOS2-AIC) except IL-6, which suggests that apart from H-bond, salt bridges and л-cation interactions, may contribute to overall stability of these complexes. Studies have reported that salt bridges and л-cation interactions are non-covalent form of interactions, which might exhibit more strong interactions than H-bond [32]. Interestingly, TMZ along with MTIC and AIC participated in salt bridge and л-cation interactions, in most of the targets, with an exception to FU. This could be attributed to the presence of L-fucose and sulphate ester groups in FU, which might attribute to its cytotoxic and anti-cancer activity[33], whereas TMZ and the metabolites possess imidazole and triazene rings, that enables formation of other non-covalent form of interactions, which contribute to conformational changes induced in the inflammatory targets. However, the exact mechanism of induction of apoptosis is still yet to be studied in detail in *in vivo* experimental models as well.

Nevertheless, there are few limitations in the current study. Firstly, the glioma cell line employed in this study is not resistant to TMZ. Thereof, expansion of this work in TMZ resistant cell lines is warranted. Secondly, effectiveness of the proposed combinatorial treatment in an *in vivo* setup is prerequisite to strengthen our *in vitro* and *in silico* findings. Also, the effect of these drugs on the anti-apoptotic markers and other pro-apoptotic signalling cascades are to be explored in mere future.

## 5.0 Conclusion

The key to accomplish successful drug repurposing is a strategy to discover the most effective drug. Here, we have demonstrated the efficacy of a marine derivative, FU on enhancing the anti-glioma potency of TMZ both in an *in vitro* and *in silico* setup. Our results, on a nutshell, indicated that the combinatorial treatment of TMZ+FU had synergistically inhibited glioma proliferation via modulation of inflammatory pathway and paving way for apoptosis. Hence, we speculate that FU could serve as a potential candidate for conventional glioma treatment enhancement.

## Acknowledgment

We would like to thank Prof C Adithan for providing “Dr Vany Adithan Research Fellowship”, centralized instrumentation, Sri Balaji Vidyapeeth (Deemed to be University) for providing the basic research infrastructure facility.

## Author Contributions

The authors declare that all data were generated in-house and that no paper mill was used. **Anitha T.S:** Conceptualization, Resources, Methodology, Supervision, Writing-Review and Editing. **Indrani Biswas:** Data Curation, Experimentation, Writing-Original draft preparation. **Daisy Precilla S** & **Shreyas S Kuduvalli**: Data Curation. **Muralidharan Arumugam Ramachandran**-Data visualization, **Akshaya S**-Data Curation, **Venkat Raman-** Data Curation, **Dhamodharan Prabhu**-Data visualization. All authors reviewed the manuscript.

## Statements and Declarations Compliance with Ethical Standards

None

---

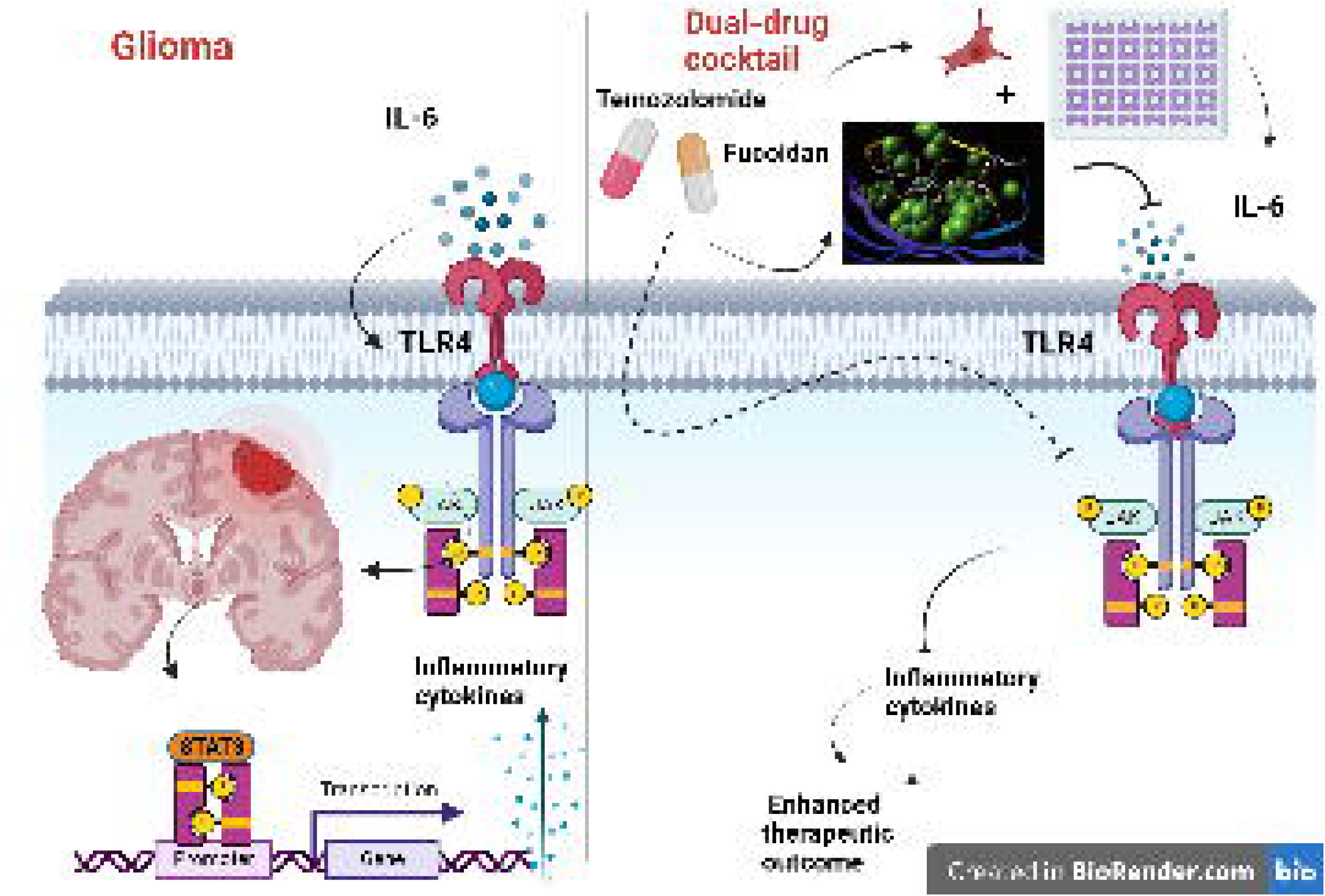

