## Supplementary Fig 1 for "Unveiling the anti-glioma potential of a marine derivative, Fucoidan: its synergistic cytotoxicity with Temozolomide-an *in vitro* and *in silico* experimental study"

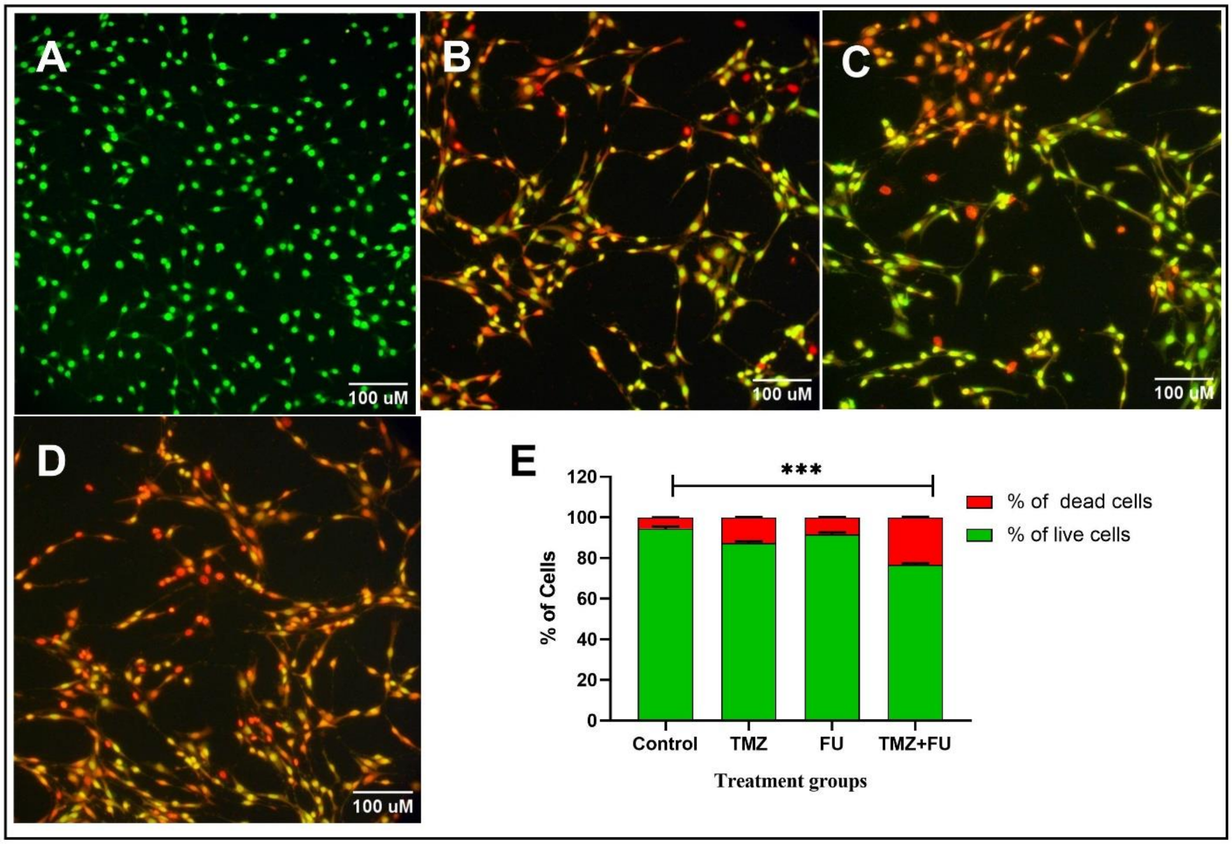


**Supplementary Fig 1:** Apoptotic cell death was observed by AO/EtBr staining under different experimental treatment groups on C6 glioma cells- (A) Control, (B) TMZ (C) FU (D) TMZ+ FU, (E) Percentage of apoptotic cells were measured by scoring viable and dead cells. *** indicates $p<0.001$ in comparison to untreated control**.**
