## Supplementary Fig 2 for "Unveiling the anti-glioma potential of a marine derivative, Fucoidan: its synergistic cytotoxicity with Temozolomide-an *in vitro* and *in silico* experimental study"

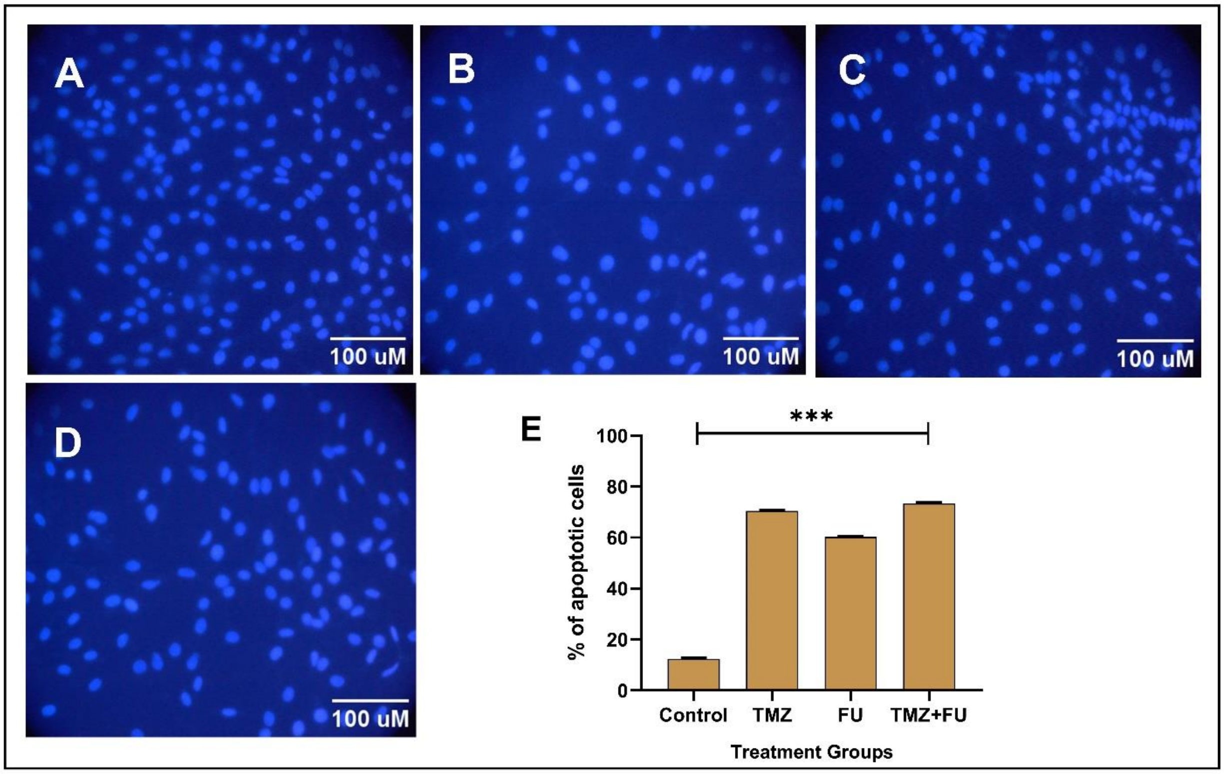


**Supplementary Fig 2:** DAPI staining under different experimental treatment groups on C6 glioma cells- (A) Control, (B) TMZ (C) FU (D) TMZ+ FU, *** indicates p<0.001 in comparison to untreated control.
