## Supplementary figures and images for "Unveiling the anti-glioma potential of a marine derivative, Fucoidan: its synergistic cytotoxicity with Temozolomide-an *in vitro* and *in silico* experimental study"

### Supplementary Fig 3

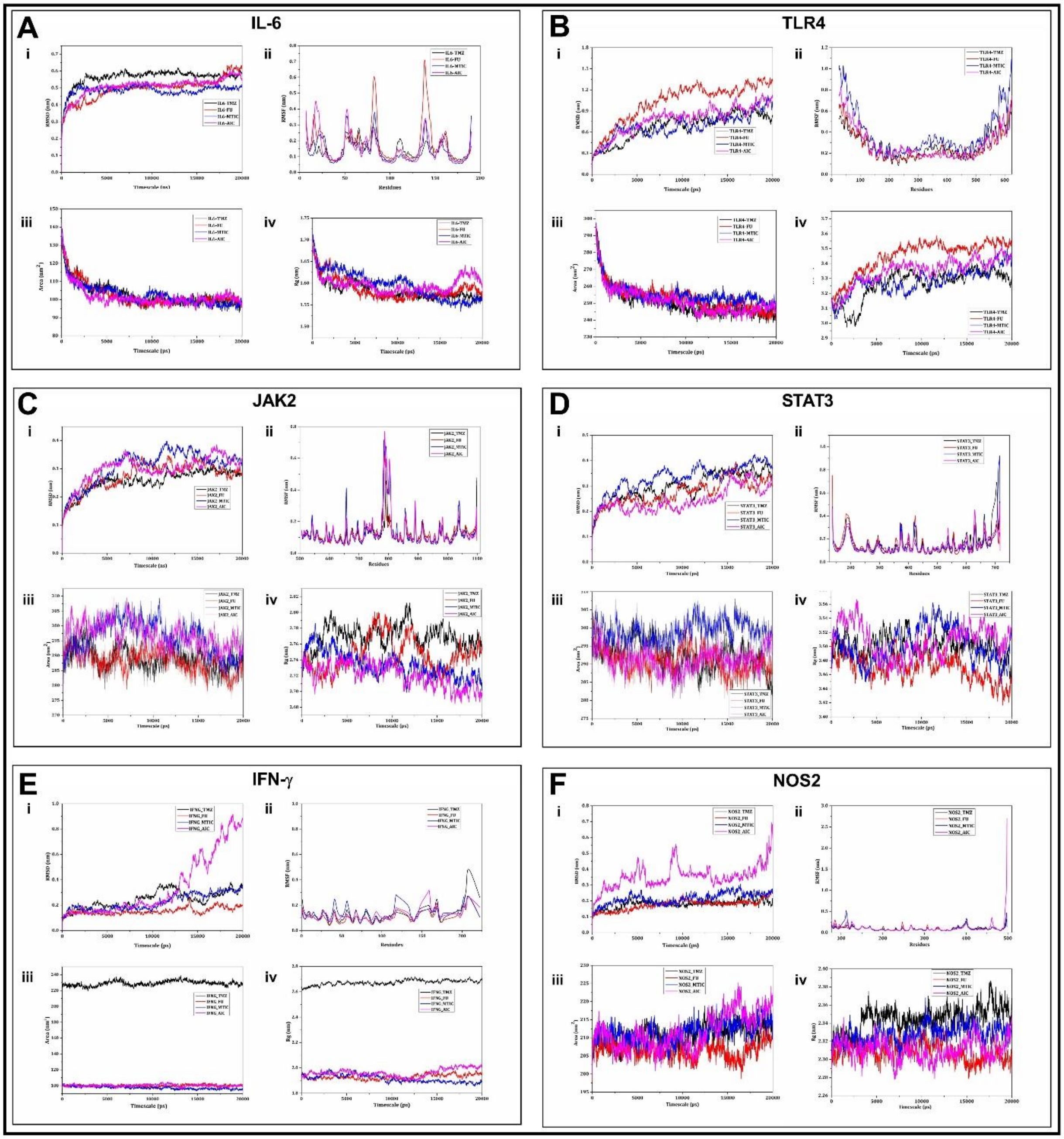


**Supplementary Fig 3:** RMSD, RMSF, SASA and Rg plots for (A) IL-6, (B) TLR4, (C) JAK2, (D) STAT3, (E) IFN-γ and (F) NOS2.
