## Supplementary Table 1 for "Unveiling the anti-glioma potential of a marine derivative, Fucoidan: its synergistic cytotoxicity with Temozolomide-an *in vitro* and *in silico* experimental study"

**Supplementary Table 1**: List of active site residues for each of the inflammatory targets

| **Sl. No.** | **Protein Targets** | **Residues involved in active site** |
|---|---|---|
| 1. | IL-6 | Ile72, Arg74, Leu89, Leu90,Lys91,Ile92,Phe179,Thr183,Ser 186 |
| 2. | TLR4 | Arg233, Phe262, Lys263, Asp264, Tyr291, Ala314, Gly315, Arg337 |
| 3. | JAK2 | Leu824, Val832, Ala849, Val880, Met898, Glu899, Tyr900, Leu901, Pro902, Glu904, Leu952 |
| 4. | STAT3 | Trp243, Lys244, Gln247, Val322, Val323, Glu324, Gln326, Thr456, His457, Pro487 |
| 5. | IFN-γ | Gln1, Met45, Gln46, Ser47, Phe60, Thr96, Asn97, Tyr98, Ser99, Val100 |
| 6. | NOS2 | Thr184, Trp188, Ala191, Cys194, Leu203, Ala237, Ile238, Phe363, Asn364, Tyr483. |
